# Competing chaperone pathways in α-synuclein disaggregation and aggregation dynamics

**DOI:** 10.1101/2025.06.25.661627

**Authors:** Nicola K. Auld, Shannon McMahon, Nicholas R. Marzano, Antoine M. van Oijen, Heath Ecroyd

## Abstract

The aggregation of the protein α-synuclein into amyloid fibrils and their subsequent deposition into large proteinaceous inclusions is a pathological hallmark of several neurodegenerative diseases, including Parkinson’s disease. Molecular chaperones, including the small heat shock proteins (sHsps) and the Hsp70 chaperone system, are known to interact with α-synuclein fibrils, preventing further aggregation and disaggregating fibrillar species respectively. However, it remains unclear if sHsps co-operate with the Hsp70 chaperones to potentially improve the kinetics or effectiveness of Hsp70-mediated disaggregation and how disaggregation kinetics are influenced by aggregation-prone α-synuclein monomers. Using thioflavin-T assays, we demonstrate that the sHsps Hsp27 (HSPB1) and αB-crystallin (HSPB5) do not synergise with the Hsp70 chaperones during α-synuclein seed fibril disaggregation. Moreover, the addition of monomeric α-synuclein with fibril seeds results in increased aggregation that overwhelms Hsp70-mediated disaggregation. Upon addition of sHsps to this system, antagonism between the two chaperone classes is observed, likely due to these chaperones competing for productive binding to the ends of α-synuclein fibrils. Overall, these results suggest that while Hsp70 and sHsp chaperones are independently capable of binding to and inhibiting fibril elongation, they do not have synergistic effects on disaggregation. Furthermore, Hsp70-mediated disaggregation is ineffectual in the presence of physiological concentrations of α-synuclein monomers, conditions that actually lead to further α-synuclein aggregation. Overall, these data may offer insight into factors that lead to the failure of the Hsp70 chaperones to clear cells of α-synuclein aggregates that leads to neurodegenerative disease.

## 1 INTRODUCTION

Many neurodegenerative diseases are associated with the formation and accumulation of stable and highly ordered filamentous protein aggregates known as amyloid fibrils, the deposition of which into insoluble intracellular or extracellular protein inclusions serve as pathological hallmarks of neurodegenerative diseases (Aguzzi and O’Connor 2010, Chiti and Dobson 2017). More specifically, the aggregation and deposition of the protein α-synuclein into Lewy body inclusions in the brain is a characteristic feature of Parkinson’s disease (PD), multiple system atrophy (MSA) and dementia with Lewy bodies (Spillantini et al. 1997), collectively known as synucleinopathies. The formation of α-synuclein fibrils from monomers occurs through a nucleation-dependent polymerisation process (Wood et al. 1999). During the initial lag phase, monomeric α-synuclein molecules associate with each other to form oligomers or pre-fibrillar species, with a characteristic cross-β-sheet structure (Sunde et al. 1997). These nuclei or “seeds” form the basis for elongation into fibrils, which proceeds rapidly through addition of monomers to the ends of these nuclei until the system reaches an equilibrium. Secondary nucleation events, including fibril breakage or fragmentation and nucleation of monomers on the surface of existing fibrils, can accelerate the aggregation process (Knowles et al. 2009, Cohen et al. 2013, Buell et al. 2014, Gaspar et al. 2017). Whilst mature fibrils, seeds and early oligomers of α-synuclein are cytotoxic, the widely accepted consensus is that oligomeric and seed forms of α-synuclein are more cytotoxic than mature fibrils (reviewed in (Alam et al. 2019, Cascella et al. 2022)).

It is well recognised that molecular chaperones are a critical first line of defence against protein aggregation (Hartl et al. 2011, Chiti and Dobson 2017). However, more recent work has demonstrated that some molecular chaperones are also involved in the disassembly of aggregates, including amyloid fibrils (Glover and Lindquist 1998, Shorter 2011, Rampelt et al. 2012, Nillegoda et al. 2015, Wentink et al. 2020). In particular, purified components of the Hsp70 chaperone system, which consists of an Hsp70 chaperone together with an Hsp40 co-chaperone and a nucleotide exchange factor (NEF), are capable of ATP-dependent disaggregation of fibrils formed by α-synuclein (Gao et al. 2015, Wentink et al. 2020, Franco et al. 2021a, Schneider et al. 2021, Beton et al. 2022), tau (Nachman et al. 2020) and huntingtin (Scior et al. 2018). A model of Hsp70-mediated disaggregation of α-synuclein fibrils suggests that the binding of DNAJB1 (an Hsp40) to the fibril promotes the recruitment and clustering of HspA8 (a constitutively expressed Hsp70; Hsc70) at the ends of fibrils (Beton et al. 2022), a process facilitated by the NEF Hsp110 (also referred to as HspH2, HspA4 and Apg-2). The association of this high molecular mass chaperone complex generates an entropic pulling force that allows the disassembly of α-synuclein fibrils (Wentink et al. 2020, Franco et al. 2021a, Schneider et al. 2021, Beton et al. 2022). There remains conjecture as to how this process occurs with some suggesting a rapid ‘unzipping’ of protofilaments generating monomeric products (Franco et al. 2021a, Schneider et al. 2021, Beton et al. 2022), whilst others report that potentially toxic oligomers and small fragments may also be generated (Gao et al. 2015, Tittelmeier et al. 2020, Beton et al. 2022, Jäger et al. 2024). These studies have led to the proposal that the Hsp70 machinery is a potential therapeutic target for neurodegenerative diseases. However, in the context of disease, Hsp70-mediated disaggregation would occur in cells in the presence of free monomeric α-synuclein, which is estimated to constitute up to 1% of cytosolic neuronal protein (Iwai et al. 1995). It is therefore important to assess the efficiency of chaperone-mediated disaggregation of α-synuclein fibrils in the presence of free monomeric α-synuclein, as free monomers in solution can potentially enhance the kinetics of aggregation. In addition, it remains to be explored whether other molecular chaperones impact the disaggregation activity of the Hsp70 chaperone system.

The small heat-shock proteins (sHsps) are a class of molecular chaperones known to inhibit the fibrillar aggregation of α-synuclein through interactions with both monomeric and oligomeric species (Rekas et al. 2004, Waudby et al. 2010, Bruinsma et al. 2011, Cox et al. 2014). Moreover, sHsps can form stable complexes with mature α-synuclein fibrils (Waudby et al. 2010, Cox et al. 2016, Selig et al. 2020). There are reports that the binding of sHsps to amyloid fibrils can lead to disaggregation; for example, the sHsp αB-crystallin (HSPB5) dissociates oligomeric forms of the amyloid fibril forming protein β_2_ microglobulin into monomers (Esposito et al. 2013, Stepanenko et al. 2020) and both Hsp27 (HSPB1) and αB-crystallin are capable of disaggregating amyloid fibrils of apolipoprotein-C (Binger et al. 2013, Selig et al. 2020). A previous study demonstrated that neither αB-crystallin nor Hsp27 alone were found to be capable of disaggregating α-synuclein fibrils (Selig et al. 2020). However, Duennwald *et al*. (2012) reported that αB-crystallin enhanced the disaggregation of α-synuclein fibrils by the Hsp70 chaperone system, albeit this work involved an (atypical) 10-day incubation of fibrils with chaperones to observe disaggregation, which is a much slower rate of disaggregation than typically observed for the Hsp70 chaperone system alone (Wentink et al. 2020, Franco et al. 2021a, Schneider et al. 2021). Hence, the capacity for sHsps to enhance disaggregation by the Hsp70 chaperone system remains unclear and warrants further investigation.

Here, we investigate whether the sHsps αB-crystallin or Hsp27 impact the Hsp70-mediated disaggregation of fibrillar forms of α-synuclein. We find no evidence for these sHsps having a synergistic role in α-synuclein fibril disaggregation by the Hsp70 chaperone system. Instead, we found an antagonistic relationship between these two chaperone classes during fibril disaggregation, likely due to the chaperones competing for binding to the ends of the fibrils. Furthermore, we find that, at physiologically relevant concentrations, the presence of α-synuclein monomers overwhelms the disaggregation capacity of the Hsp70 chaperones, resulting in overall elongation of fibrils.

## 2 RESULTS

### 2.1 The Hsp70 chaperone system removes α-synuclein monomers from fibril ends

Hsp70-mediated disaggregation of fibrillar forms of α-synuclein is hypothesised to progress rapidly from fibril ends generating monomeric products (Franco et al. 2021a) but may also include fibril fragmentation (Gao et al. 2015, Tittelmeier et al. 2020, Beton et al. 2022). To investigate this, recombinant α-synuclein was aggregated to form mature fibrils, which were subsequently sonicated to produce small amyloid fragments (seeds). As expected, seeds and mature fibrils of α-synuclein were successfully disaggregated by the Hsp70 system chaperone machinery (Figure 1A) in an ATP-dependent manner (Supplementary Figure 1); however, the rate and amount of disaggregation was significantly greater for the α-synuclein seeds (Figure 1C, t_2.99_ = 10.61, P = 0.00182), in agreement with previous results (Franco et al. 2021a). TIRF microscopy images (Figure 1B) confirmed that seed fragments were shorter than mature fibrils (Supplementary Figure 2) and represent an increase in the number of fibril ends.

**Figure 1:**
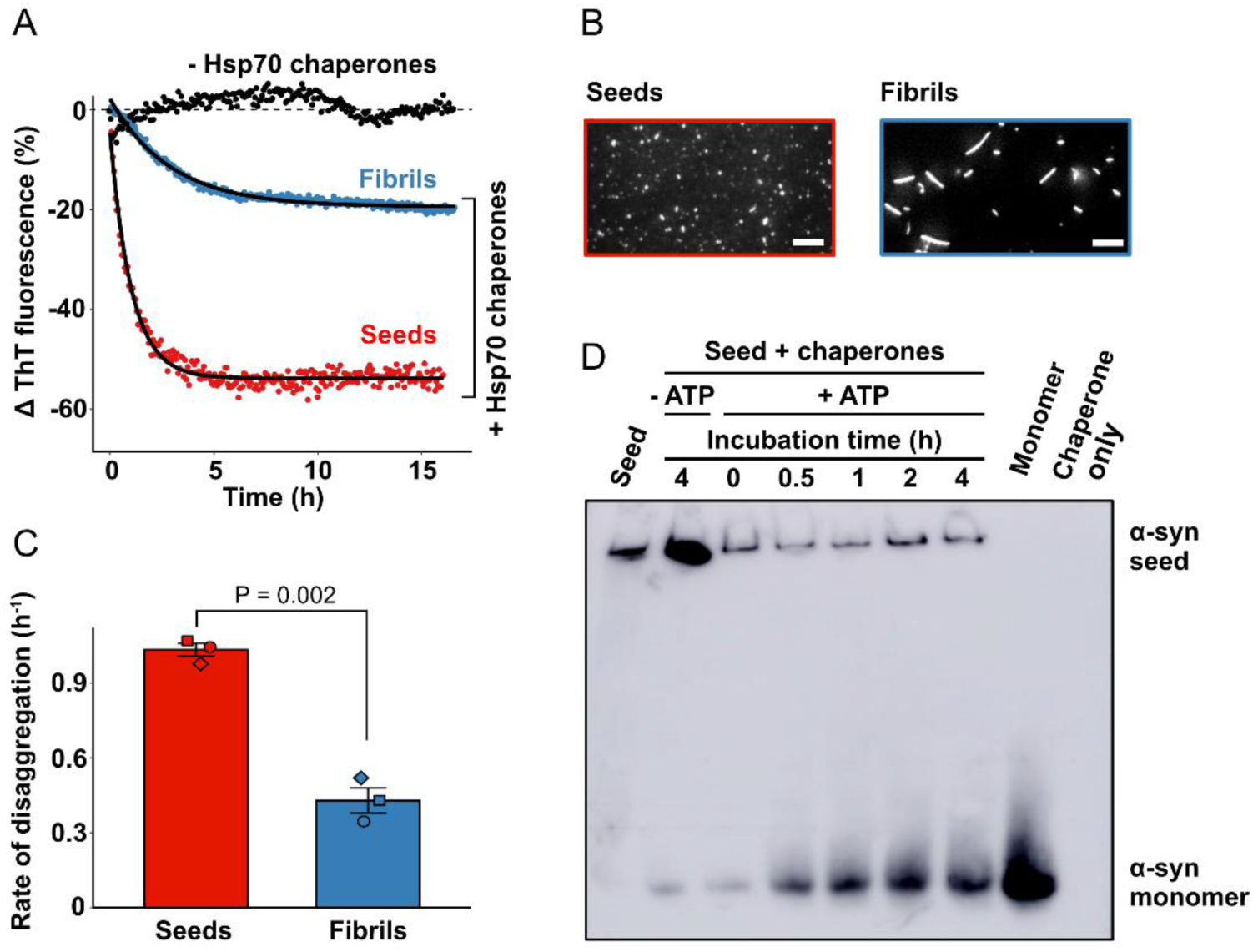
Disaggregation of α-synuclein seeds and fibrils by the Hsp70 chaperone system results in the release of free α-synuclein monomers. α-synuclein seeds or fibrils (2 µM) were incubated in the presence or absence of the Hsp70 chaperones (2 µM HspA8, 1 µM DNAJB1 and 0.2 µM Hsp110) at 30°C for up to 8 h. (A) Kinetic traces of α-synuclein seed and fibril disaggregation as monitored by the change in ThT fluorescence over time. Data is representative of three independent experiments and fitted to a one-phase exponential decay model. (B) Representative TIRF microscopy image of α-synuclein seeds and fibrils incubated in 50 mM phosphate-TROLOX buffer supplemented with 5 µM ASCP dye. Scale bars represent 5 µm. (C) The rate of seed and fibril disaggregation as determined from one-phase exponential decay model. Data are means (± SEM) of three independent experiments (n = 3) and were analysed using a student’s t-test. (D) Aliquots from the disaggregation reaction of α-synuclein seeds were taken at the timepoints indicated and added to 50 mM EDTA to quench the reaction. Samples were subject to native-PAGE and immunoblotting with an anti-α-synuclein antibody. A sample of monomeric α-synuclein (5 µM) was also included. The blot shown is representative of three independent experiments.

To investigate the products of Hsp70-mediated disaggregation, aliquots taken during the disaggregation reaction were quenched with the addition of 50 mM EDTA, to chelate MgCl_2_ and stop ATP hydrolysis by HspA8, and analysed by native-PAGE electrophoresis and subsequent immunoblotting (Figure 1D). Incubation of α-synuclein seeds with the Hsp70 chaperone system and ATP resulted in a decrease in intensity of the higher molecular weight species, indicative of a decrease in the amount of seed. Correspondingly, there was an increase in intensity of a low molecular weight band corresponding to monomeric α-synuclein. Of note, no other oligomeric forms of α-synuclein (e.g. dimers or other multimers) were detected during the disaggregation process. Thus, these data indicate that the Hsp70-mediated disaggregation of α-synuclein seeds primarily occurs through the liberation of monomeric units from fibril ends.

### 2.2 Hsp70-mediated disaggregation is outcompeted by aggregation-prone monomeric α-synuclein

Given that in the cellular environment disaggregation of α-synuclein fibrillar species occurs in the presence of monomeric α-synuclein, we investigated the capacity of the Hsp70 chaperones to disaggregate α-synuclein seed fibrils when additional free monomeric α-synuclein was present. As expected, there was a concentration-dependent increase in the overall change in ThT fluorescence associated with fibril elongation in the presence of higher concentrations of monomeric α-synuclein (Figure 2A). Interestingly, only a relatively modest concentration of α-synuclein monomers (2 µM) negated the decrease in ThT fluorescence associated with Hsp70-mediated disaggregation of α-synuclein seed fibrils, shifting the overall kinetics towards aggregation, as evidenced through a net increase in ThT fluorescence (Figure 2B). When the Hsp70 chaperones were present, the overall level of fibril aggregation stimulated by monomeric α-synuclein was reduced by up to ∼3-fold (Figure 2C). Furthermore, the constitutively expressed HspA8 and its cochaperone DNAJB1 were responsible for the inhibition of fibril elongation, offering significantly more protection against seed elongation than the nucleotide exchange factor Hsp110 and the non-chaperone control protein SOD1 (Figure 2D, F_4,48_ = 108, P < 0.0001). Together, these data suggest that in the presence of free monomeric α-synuclein, the Hsp70 chaperone system can suppress the amount of fibrillar aggregation that would otherwise occur, but is unable to effectively promote fibril disaggregation.

**Figure 2:**
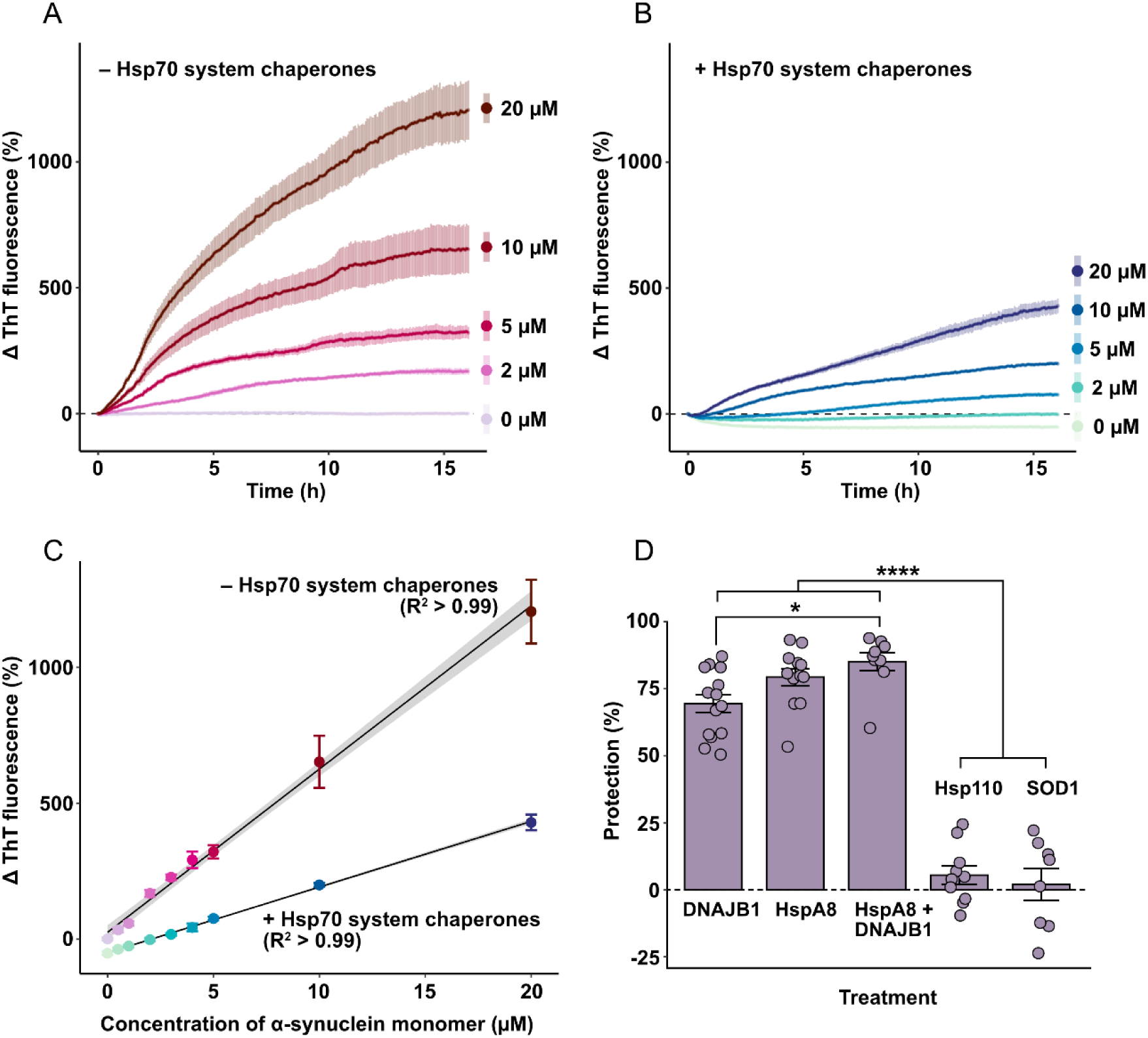
The presence of free monomeric α-synuclein reduces the capacity of the Hsp70 chaperone system to mediate disaggregation and shifts the reaction towards fibril aggregation. α-synuclein seeds (2 µM) were incubated with increasing concentrations of monomeric α-synuclein (0 – 20 µM) in the (**A**) absence or **(B)** presence of the Hsp70 chaperones (2 µM HspA8, 1 µM DNAJB1 and 0.2 µM Hsp110) in disaggregation buffer containing 20 µM ThT at 30 °C. The change in ThT fluorescence was monitored over time. Data are means (± SEM) of 3 independent experiments (n = 3). (**C**) The end-point normalised ThT fluorescence for each treatment as a function of the concentration of free monomeric α-synuclein present. Data were fit to a linear regression model. **(D)** α-synuclein seeds (2 µM) were incubated with 10 µM monomeric α-synuclein and individual components of the Hsp70 chaperone system (or an equivalent amount of the non-chaperone control protein SOD1) in disaggregation buffer containing 20 µM ThT at 30°C. The percentage protection against seed elongation afforded by each chaperone combination is shown. Data are means (± SEM) of at least 8 independent experiments (n ≥ 8) and analysed using a one-way ANOVA with Tukey’s HSD post-hoc testing where * and **** indicate P < 0.05 and P ≤ 0.0001, respectively (comparisons not indicated were not significant; P > 0.05).

### 2.3 sHsps antagonise Hsp70 chaperones during disaggregation

Given that sHsps are known to bind to α-synuclein fibrils (Waudby et al. 2010, Cox et al. 2016) and have some capacity to disaggregate fibrils formed by apolipoprotein C-II (Selig et al. 2020) or β_2_ microglobulin (Esposito et al. 2013), we first examined whether the presence of sHsps alone resulted in a decrease in the ThT fluorescence associated with α-synuclein seed fibrils. No decrease in ThT fluorescence was observed when α-synuclein seeds were incubated with sHsps alone at any of the concentrations tested (Figure 3A). This result is in agreement with previous data that indicated that neither αB-crystallin nor Hsp27 were capable of causing appreciable levels of disaggregation of fibrillar forms of α-synuclein in isolation (Selig et al. 2020).

**Figure 3:**
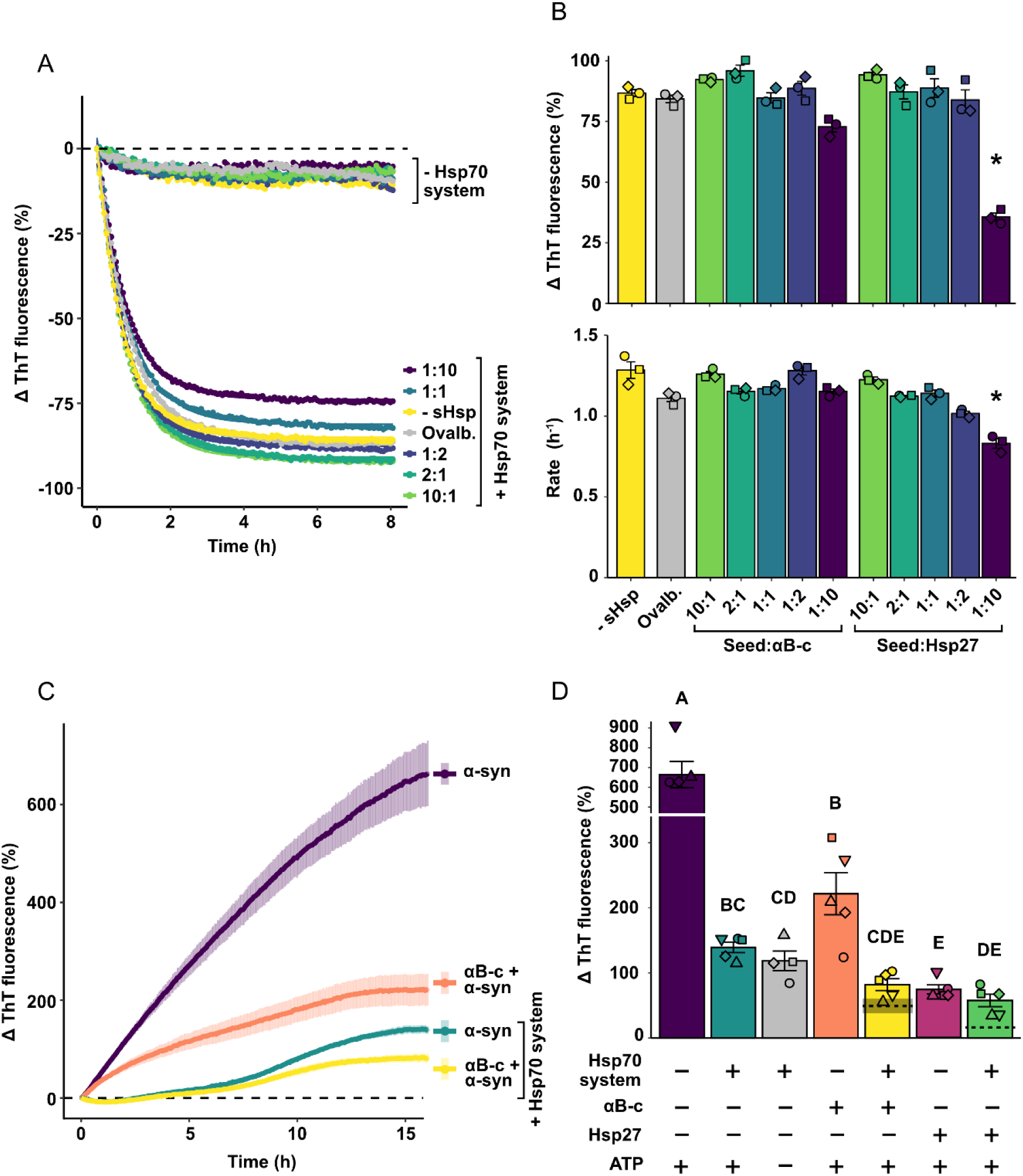
sHsps do not cooperate with Hsp70 system chaperones during disaggregation but act antagonistically to prevent α-synuclein seed fibril elongation. **(A)** A representative trace of the α-synuclein seeds pre-incubated with sHsps, with and without the Hsp70 chaperone system. Data are from technical duplicates of each sample, and those with the Hsp70 chaperones were fitted to a one-phase exponential decay model from which the **(B)** rate and amount of disaggregation were determined. Statistically significant differences between the means of all sHsp concentrations, ovalbumin, and the seed only control were assessed using a one-way ANOVA and Tukey’s HSD post-hoc test for the rate of disaggregation (P < 0.0001) and the overall amount of disaggregation (P < 0.0001). Treatments that were significantly different from both the seed only (-sHsp) and non-chaperone (ovalbumin) controls are indicated by *. **(C)** Representative traces showing α-synuclein seeds incubated with combinations of sHsps, the Hsp70 system and ATP in addition to 10 µM monomeric α-synuclein in disaggregation buffer containing 20 µM ThT at 30°C for 16 h. Data are means (± SEM) of at least 4 independent experiments. **(D)** The overall normalised change in ThT fluorescence (as a percentage) was analysed via a one-way ANOVA (P < 0.0001) with a Tukey’s HSD post-hoc test with significant differences between treatments indicated via differing letters. Expected additive effects of the combination of each sHsp with the Hsp70 chaperones are indicated as dashed lines (mean ± SEM), with SEM denoted by shading. In some cases, SEM is too small to be visualised).

Next, we explored whether sHsps influence the Hsp70-mediated disaggregation of α-synuclein seeds. To facilitate sHsp binding, the seeds were pre-incubated with either αB-crystallin or Hsp27, at varying molar ratios (from 10:1 – 1:10; α-synuclein seed:sHsp). Upon dilution of sHsp-bound seeds into buffer containing the Hsp70 system chaperones, the characteristic exponential decay in ThT fluorescence was observed, indicative of disaggregation (Figure 3A). Analysis by one-way ANOVA (Figure 3B, F_11,24_ = 43.2, P < 0.0001) and subsequent post-hoc testing demonstrated that at low molar ratios (10:1, 2:1, 1:1 and 1:2 seed:sHsp) there was no significant difference in the amount of disaggregation that occurred in the presence or absence of the sHsps (P ≥ 0.0828). At the highest molar ratio tested (1:10 seed:sHsp), the addition of Hsp27 resulted in significantly lower levels of seed disaggregation compared to when the sHsp was not present or when the non-chaperone control protein ovalbumin was present (P < 0.0001). With regard to the rate at which disaggregation occurs, again only the highest concentration of Hsp27 (a molar ratio of 1:10 seed:Hsp27) significantly decreased the rate of disaggregation compared to both the absence of the sHsp or when ovalbumin was present (P < 0.0001). This suggests that high concentrations of Hsp27 relative to the amount of α-synuclein seed decreases both the rate of Hsp70-mediated disaggregation and the amount of disaggregation that occurs.

Since sHsps are potent suppressors of α-synuclein fibrillar aggregation (Bruinsma et al. 2011, Cox et al. 2016), we next investigated whether these sHsps further reduce the net elongation of α-synuclein seeds observed in samples containing seeds, the Hsp70 chaperone system and free monomeric α-synuclein. Whilst the combination of αB-crystallin and the Hsp70 chaperones significantly reduced the elongation of fibrils compared to when αB-crystallin was present alone (F_6,27_ = 46.6, P < 0.0001), there was no significant difference compared to when the Hsp70 chaperones were present alone (Figure 3C-D). Similar results were obtained when Hsp27 was present in place of αB-crystallin. The expected change in ThT fluorescence arising from the addition of each sHsp with the Hsp70 chaperones, in the presence of both seeds and free monomer was calculated (indicated by dashed lines in Figure 3D). As neither Hsp27 nor αB-crystallin in combination with the Hsp70 chaperones achieved the level of aggregation suppression that would be indicative of synergy or independent action, these data indicate that sHsps and Hsp70 chaperones act antagonistically to inhibit the elongation of α-synuclein seeds by monomeric α-synuclein.

### 2.4 Hsp70-mediated disaggregation promotes α-synuclein aggregation in the presence of sHsps at physiologically relevant concentrations

Given the interplay between monomeric and fibrillar forms of α-synuclein when chaperones are present, we sought to examine the impact of physiologically relevant concentrations of free monomeric α-synuclein and the molecular chaperones with regard to the aggregated state of α-synuclein. To do so, we used concentrations of α-synuclein and the chaperones based on previously reported values: namely 50 µM monomeric α-synuclein (Iwai et al. 1995, Wilhelm et al. 2014); 20 µM sHsps (Mymrikov et al. 2020); and 14 µM HspA8 (Moran Luengo et al. 2018) with concentrations of DNAJB1 and Hsp110 so as to maintain the 1:0.5:0.1 (HspA8:DNAJB1:Hsp110) molar ratio used previously for the Hsp70 chaperone system. Of note, when seed fibrils were incubated with physiologically relevant concentrations of free monomeric α-synuclein there were significant increases in ThT fluorescence (F_7,48_ = 62.4, P < 0.0001) even when the Hsp70 chaperone disaggregation machinery was present (Figure 4).

**Figure 4.**
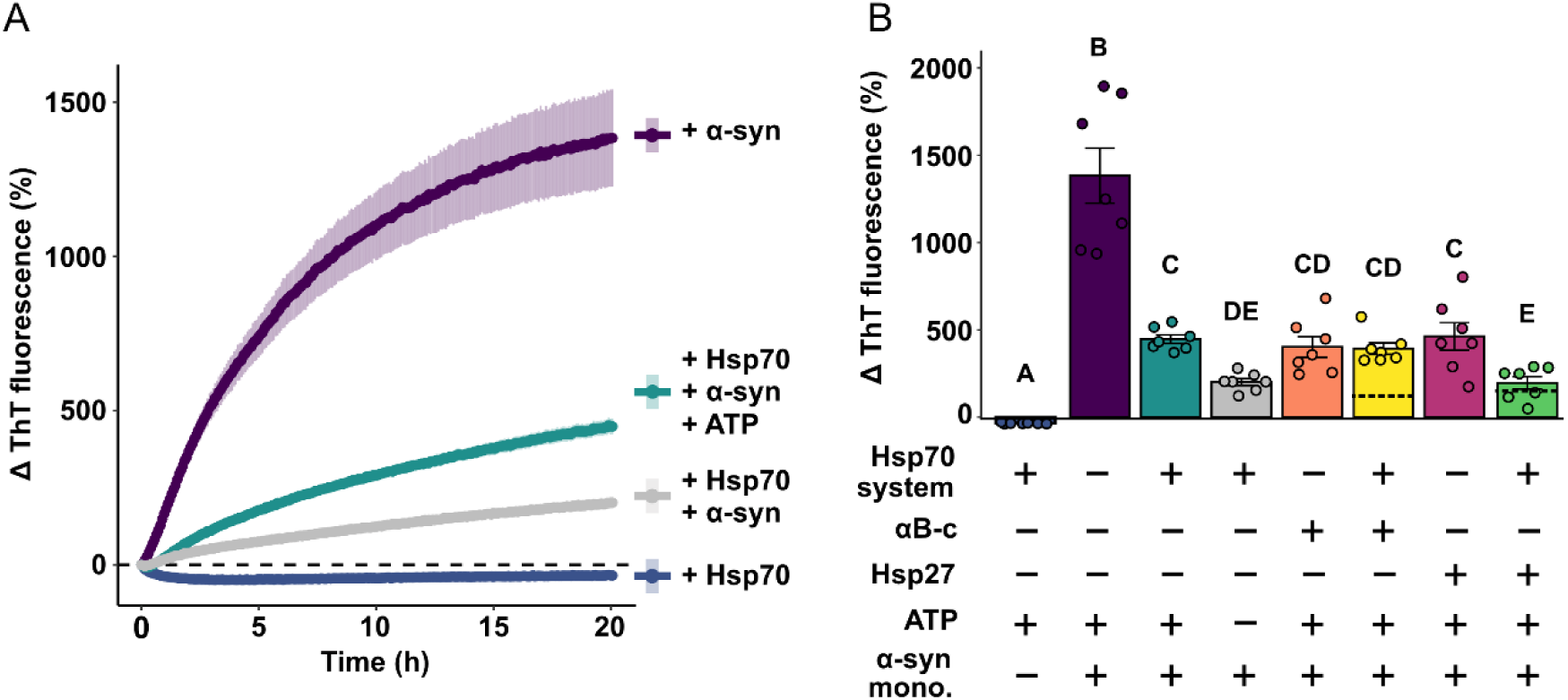
The disaggregation of α-synuclein seeds at physiological concentrations of Hsp70 and sHsps. Physiologically relevant concentrations of Hsp70 (14 µM) and Hsp27 (20 µM) were incubated with α-synuclein seeds (2 µM) and free monomeric α-synuclein (50 µM). Select traces are shown in **(A)** where data are means (± SEM) of 7 independent experiments. The overall normalised change in ThT fluorescence (as a percentage) **(B)** was analysed via a one-way ANOVA (P < 0.0001) with a Tukey’s HSD post-hoc test with significant differences between treatments indicated via differing letters. Expected additive effects of the combination of each sHsp with the Hsp70 chaperones are indicated as dashed lines (mean ± SEM), with SEM denoted by shading. In some cases, SEM is too small to be visualised.

Strikingly, the presence of ATP with the Hsp70 chaperones led to a significant increase in ThT fluorescence compared to when ATP was absent (Figure 4B, P = 0.0105), suggesting that active Hsp70-mediated disaggregation enhances α-synuclein aggregation under these conditions. In line with previous findings (see Figure 3C-D), no evidence of synergy between chaperone classes was observed. Specifically, the combination of αB-crystallin and the Hsp70 chaperones displayed evidence of antagonism and the combination of Hsp27 and the Hsp70 chaperones was observed to be independent or simply additive (Figure 4B).

## 3 DISCUSSION

The chaperone network plays a key role in regulating the aggregation of α-synuclein, a process that, when left unchecked, leads to the formation of amyloid fibrils, which are pathological hallmarks of the synucleinopathies. Of therapeutic interest are chaperones that interact with fibrillar forms of α-synuclein, potentially leading to their disaggregation. The ATP-dependent Hsp70 system chaperones constitute a disaggregation machinery that is capable of disaggregating α-synuclein fibrils (Gao et al. 2015, Wentink et al. 2020) whereas the ATP-independent sHsps bind directly to α-synuclein fibrils, preventing their further elongation, but are not independently capable of mediating disaggregation (Selig et al. 2020). However, it remained to be established whether the binding of sHsps to α-synuclein fibrils impacts Hsp70-mediated amyloid fibril disaggregation. Here, we provide evidence that these two functionally distinct classes of molecular chaperones have antagonistic, rather than synergistic, effects when it comes to mediating α-synuclein fibril disaggregation. Furthermore, we demonstrate that the presence of free monomeric α-synuclein significantly reduces the efficacy of Hsp70 chaperones to disassemble α-synuclein fibrils *in vitro*.

The overwhelming majority of studies investigating chaperone-mediated α-synuclein fibril disaggregation involve the study of exclusively fibrillar forms of α-synuclein. However, in a physiological setting within the cell, α-synuclein aggregates form and elongate within a pool of monomeric α-synuclein. Mouse models of synucleinopathies reveal a prion-like spreading and amplification of fibrillar α-synuclein following inoculation of mice with pre-formed fibrils (Luk et al. 2012, Masuda-Suzukake et al. 2013, Ayers et al. 2018), highlighting the role that monomeric α-synuclein plays in mediating the spread of disease-related pathology. Hence, we sought to investigate the kinetics of Hsp70-mediated α-synuclein seed disaggregation in the presence of free monomeric α-synuclein. Our data show that, in samples containing Hsp70 chaperones and fibrillar forms of α-synuclein, as the amount of free monomeric α-synuclein increases, the ThT fluorescence kinetics are characterised by an increasingly brief disaggregation phase followed by elongation of the fibrils (see Figure 2B). Moreover, the addition of a relatively low (2 µM) amount of free monomeric α-synuclein was sufficient to negate the overall reduction in ThT fluorescence associated with the disaggregation facilitated by the Hsp70 chaperones. The decrease in disaggregation observed in the presence of α-synuclein monomers may be due to competition between monomeric and aggregated forms of α-synuclein for Hsp70 chaperone binding, whereby increased levels of monomers in solution act to sequester the Hsp70 chaperones from binding to fibrils. However, Hsp70 and DNAJ isoforms related to those used in this study have been demonstrated to preferentially bind to fibrillar, as opposed to monomeric, forms of α-synuclein (Hinault et al. 2010, Aprile et al. 2017). Alternatively, the kinetic and thermodynamic advantages of the association of monomers with fibrils may surpass those of chaperone-mediated disaggregation, resulting in the predominance of seed elongation over disaggregation. Regardless, the relatively modest concentration of α-synuclein required to negate the effects of Hsp70 chaperone-mediated disaggregation suggests that a delicate balance exists between the competing forces of seed elongation and disaggregation in the cellular environment.

Whilst disaggregation was limited in the presence of free α-synuclein monomers, the presence of Hsp70 chaperones under these conditions did suppress the increase in ThT fluorescence associated with seed elongation (see Figure 2C). This observation suggests that, under these circumstances, the Hsp70 and DNAJ chaperones can bind α-synuclein to prevent further aggregation, in agreement with previous results (McLean et al. 2002, Klucken et al. 2004, Luk et al. 2008, Pemberton et al. 2011). Together, these results highlight the ability of the Hsp70 chaperone system to not only act as a “disaggregase” but additionally in a “holdase” role to mitigate α-synuclein seed elongation in the presence of α-synuclein monomers. Furthermore, the addition of sHsps to the system containing Hsp70 chaperones, monomeric α-synuclein and seeds, resulted in no synergistic suppression of seed elongation (Figure 3C-D). Instead, the sHsp and Hsp70 chaperones were found to be antagonistic with regard to the change in ThT fluorescence associated with α-synuclein aggregation under these conditions. As both sHsps (Waudby et al. 2010, Selig et al. 2022) and the Hsp70 chaperones (Franco et al. 2021a, Schneider et al. 2021, Beton et al. 2022) are thought to bind preferentially to fibril ends, the antagonism observed may likely be due to competition for binding to these end regions of the fibrils.

We note incomplete or partial disaggregation of α-synuclein fibrils by the Hsp70 chaperone system, evidenced by the presence of fibrillar α-synuclein species at the end of the 4 h incubation period, by which time the decrease in ThT fluorescence has reached a plateau (Figure 1A and D). Similar findings have been reported in both biochemical (Gao et al. 2015) and atomic force microscopy (Beton et al. 2022) studies indicating that some fibrillar species remain unaffected by the disaggregation machinery (Beton et al. 2022). As α-synuclein is able to form a variety of polymorphic oligomeric and fibrillar structures (reviewed in (Mehra et al. 2021, Yoo et al. 2022)), with some morphologies suggested to be more amenable to chaperone-mediated disaggregation than others (Fricke et al. 2024, Jäger et al. 2024), it is possible that structural differences may account for the persistence of some fibrillar species seen in this work.

It has been demonstrated previously that truncation of α-synuclein, resulting in the removal of negative charges at the C-terminal region (Franco et al. 2021b) and the removal of key N-terminal HspA8 interaction sites at residues 1-10 and 37-43 (Redeker et al. 2012, Wentink et al. 2020), can reduce or completely abolish disaggregation by the Hsp70 system chaperones. These results highlight the need for HspA8 and its co-chaperone DNAJB1 to access specific regions of α-synuclein in order for productive binding and disaggregation of fibrils to proceed. It follows that fibrils that are not fully disaggregated may persist, proliferate and amplify – a mechanism that may lead to the accumulation of certain α-synuclein polymorphs in deposits such as Lewy bodies. Furthermore, incomplete disaggregation may lead to the generation of α-synuclein species that are more capable of prion-like spreading in a disease context (Silveira et al. 2005, Bett et al. 2017).

Whilst co-aggregation of sHsps with amorphously aggregating client proteins is crucial for efficient resolubilisation and disassembly of these aggregates by the Hsp70 system chaperones (Rampelt et al. 2012, Nillegoda et al. 2015, Gonçalves et al. 2021), only a single study has previously investigated the effect of sHsps on Hsp70-mediated amyloid disaggregation (Duennwald et al. 2012). In line with previous observations (Selig et al. 2020), we found that sHsps alone were not capable of disaggregating α-synuclein seeds to any appreciable extent. This suggests that the binding of sHsps to α-synuclein fibrils acts to stabilise the fibril, preventing further elongation and decreasing toxicity (Cox et al. 2018, Selig et al. 2020). Moreover, when sHsp-bound α-synuclein fibrils were exposed to the Hsp70 disaggregation machinery, we found no evidence to suggest a synergistic effect of these two chaperone systems on α-synuclein disaggregation. Instead, given the observation that when free α-synuclein monomer is present, the two chaperone classes are antagonistic with regard to suppressing further α-synuclein aggregation, our data suggests that sHsps and Hsp70 chaperones compete for binding to the ends of α-synuclein fibrils, since this is where monomers are removed during disaggregation.

Increasing the concentrations of sHsps, Hsp70 chaperones and monomeric α-synuclein present with α-synuclein seeds to approximate physiologically relevant levels led to an overall increase in ThT fluorescence over time. Thus, under these conditions, disaggregation activity is overwhelmed by the rate at which seeds elongate and aggregate in the presence of monomers. The observation that the presence of ATP promotes seed aggregation (Figure 4B) suggests that Hsp70 disaggregation activity may facilitate the proliferation and spread of seeding-competent species in a disease context. These results support previous findings showing that α-synuclein (Jäger et al. 2024) and tau (Nachman et al. 2020) fibrils subjected to Hsp70-mediated disaggregation had a greater seeding capacity than fibrils not subjected to Hsp70-mediated disaggregation. In addition, depletion of Hsp110 in a *Caenorhabditis elegans* model of disaggregation reduced both Hsp70-mediated disaggregation as well as aggregated α-synuclein foci and toxicity (Tittelmeier et al. 2020). Together, these data suggest that while disaggregation may reduce the length of α-synuclein fibrils and eliminate some aggregated species, it may ultimately lead to increased α-synuclein aggregation.

## 4 CONCLUSIONS

The disaggregation of α-synuclein fibrils by the Hsp70 chaperones is a complex and dynamic process. The results of this work indicate that although Hsp70 chaperones can efficiently reduce aggregate levels in isolation, the addition of monomeric α-synuclein reduces or negates this effect, with Hsp70 chaperones acting in “holdase” roles to prevent fibril elongation. Additionally, our results suggest that rather than synergising with Hsp70 chaperones during disaggregation, sHsps play an antagonistic role, likely competing with Hsp70 chaperones for productive binding sites on the ends of α-synuclein fibrils. Furthermore, at physiologically relevant concentrations, Hsp70 disaggregation activity may lead to an increase in the number of aggregates, potentially leading to a proliferation of species that can amplify synucleinopathy disease pathology.

## 5 MATERIALS AND METHODS

### 5.1 Protein expression, extraction and purification of α-synuclein, Hsp27 and αB-crystallin

A single colony of BL21(DE3) *Escherichia coli* (*E. coli*) cells transformed with a plasmid encoding either α-synuclein, Hsp27 or αB-crystallin was used to inoculate a starter culture (∼ 100 mL) of lysogeny broth (LB; 5% (w/v) yeast, 10% (w/v) NaCl, 10% (w/v) tryptone, pH 7.4) supplemented with appropriate antibiotics (100 µg/mL ampicillin or 50 µg/mL kanamycin). Starter cultures were grown overnight at 37°C with constant agitation at 180 rpm. The starter culture was then diluted 20-fold into LB media containing the appropriate antibiotic to ensure maintenance of the plasmid. Cultures were incubated at 37°C with shaking at 180 rpm until an optical density at 600 nm (OD_600_) of ∼ 0.8 was reached. The expression of α-synuclein, Hsp27 or αB-crystallin was induced by addition of 0.25 mM isopropyl-β-D-1-thiogalactopyranoside (IPTG) and cultures were incubated at 37°C for 4 h. The cells were then harvested by centrifugation at 5,000 × *g* for 10 min at 4°C and the pellet stored at −20°C until the recombinant protein was extracted.

In order to purify recombinant Hsp27 and αB-crystallin, cell pellets were thawed and resuspended in ice-cold lysis buffer (50 mM Tris, 100 mM NaCl and 1 mM EDTA, 0.5 mM PMSF, pH 8.0) containing a cOmplete^™^ EDTA-free protease inhibitor cocktail (Sigma-Aldrich, St. Louis, MO, USA), lysozyme (0.5 mg/mL) and DNase I (3.0 µg/mL). Resuspended cells were incubated for 30 min at 4°C with gentle rocking and then probe sonicated using the Sonifer^®^ 250 Digital cell disruptor (Branson Ultrasonics Corporation, Brookfield, CT, USA) for 3 min at 45% amplitude (10 s on/20 s off). The lysates were clarified by at least two rounds of centrifugation at 24,000 × *g* for 30 min at 4°C prior to being filtered through a 0.45 µm filter. The bacterial lysates were loaded onto a HiPrep^™^ DEAE FF 16/10 column (GE HealthCare, Chicago, IL, USA) pre-equilibrated in 20 mM Tris, 1 mM EDTA, 3 mM sodium azide, 0.5 mM PMSF (pH 8.5) and proteins were eluted at 2.5 mL/min over 8 column volumes using a linear gradient (0 – 200 mM NaCl). Fractions (5 mL) identified to contain Hsp27 or αB-crystallin were pooled and concentrated to 5 mL using a Amicon^®^ stirred filtration cell (Merck Millipore, Burlington, MA, USA) before being loaded onto a HiPrep^™^ 26/60 Sephacryl^®^ S-300 HR column (Cytiva, Marlborough, MA, USA) equilibrated in 50 mM phosphate buffer with 0.5 mM PMSF (pH 7.4). Proteins were eluted at 1.5 mL/min into 10 mL fractions, prior to being pooled, concentrated using a 10K MWCO Pierce^™^ Protein Concentrator (Thermo Fisher Scientific, Waltham, MA, USA) and stored at −80°C until required.

Purification of recombinant α-synuclein was performed as described previously (Buell et al. 2014) with modifications. Briefly, bacterial cells were resuspended in lysis buffer (100 mM Tris and 10 mM EDTA, pH 8.0) supplemented with protease inhibitor, prior to being subjected to three slow freeze-thaw cycles (−20°C to room temperature with gentle rocking). The lysate was then probe sonicated as described above before being heated to 90°C for 20 min to remove bacterial host proteins. The lysate was centrifuged at 22,000 × *g* for 20 min at 4°C. Streptomycin sulfate was added to the cleared lysate at a concentration of 10 mg/mL and was stirred at 4°C for 20 min. This process of centrifugation and subsequent stirring in the presence of streptomycin sulfate was repeated once before the lysate was clarified by a final round of centrifugation at 24,000 × *g* for 20 min at 4°C. The lysate was filtered through a 0.45 µm filter and applied to a HiPrep^™^ DEAE FF 16/10 column (Cytiva) equilibrated in 20 mM Tris, 1 mM EDTA, 3 mM sodium azide (pH 8.5). The protein was eluted at 2.5 mL/min over 4 column volumes using a linear gradient (0 – 500 mM NaCl) and 5 mL fractions were collected.

Fractions identified to contain α-synuclein were pooled and loaded onto a HiLoad^®^ 16/600 Superdex^®^ 75 pg size-exclusion column (GE HealthCare) pre-equilibrated in 50 mM phosphate buffer (pH 7.5) and the protein was eluted into 4 mL fractions at a flow rate of 1.0 mL/min. Fractions containing purified, recombinant α-synuclein were pooled and the protein was concentrated using a 10K MWCO Pierce^™^ Protein Concentrator prior to storage at −80°C.

### 5.2 Protein expression, extraction and purification of HspA8, DNAJB1 and Hsp110

Plasmids encoding recombinant 6×His-SUMO-HspA8 (also known as Hsc70), 6×His-SUMO-DNAJB1 (also known as Hdj-1) or 6×His-SUMO-Hsp110 (also known as HspA4, HspH2 and Apg-2) were transformed into Rosetta^™^(DE3) *E. coli* and single colonies were used to inoculate ∼ 100 mL 2× yeast extract tryptone media (YT; 10% (w/v) yeast extract, 5% (w/v) NaCl, 16% (w/v) tryptone, pH 7.0) media supplemented with kanamycin (50 µg/mL) and chloramphenicol (25 µg/mL). The starter cultures were incubated overnight at 37°C with shaking at 180 rpm before being diluted 20-fold into fresh 2×YT media supplemented with kanamycin (50 µg/mL) only. Once cultures reached an OD_600_ of between 0.6 – 0.8, expression was induced using 1 mM IPTG. Cells were grown for a further 4 (HspA8 and DNAJB1) to 6 h (Hsp110) at 30°C with constant shaking before being harvested by centrifugation at 5,000 × *g* for 10 min at 4°C. Cell pellets were stored at −20°C until extraction. Cells containing recombinant HspA8, DNAJB1 or Hsp110 were resuspended in lysis buffer (50mM HEPES-KOH (pH 7.4), 5 mM MgCl_2_, 2 mM β-mercaptoethanol (BME), 20 mM imidazole, 10% glycerol) supplemented with a protease inhibitor cocktail, DNase I (3.0 μg/ml), lysozyme (0.5 mg/mL) and either 750 mM (DNAJB1) or 300 mM (HspA8 and Hsp110) KCl. Cells were sonicated as described previously and centrifuged (24,000 × *g*, 30 min, 4°C) twice before being passed through a 0.45 μm filter.

HspA8, DNAJB1 and Hsp110 were purified by loading each bacterial cell lysate (∼ 50 mL) at 1 mL/min onto a 5 mL HisTrap^™^ HP Sepharose column (Cytiva) equilibrated with the appropriate lysis buffer. The column was then washed at 5 mL/min with a low salt buffer (50 mM HEPES-KOH (pH 7.4), 5 mM MgCl_2_, 2 mM BME, 20 mM imidazole, 10% glycerol) containing either 500 mM (DNAJB1) or 100 mM (HspA8 and Hsp110) KCl until the absorbance at 280 nm had returned to baseline. For HspA8 and Hsp110, an additional low salt buffer containing 10 mM ATP was applied to the column and the flow stopped for 30 min at room temperature. The column was washed again with low salt buffer until the absorbance at 280 nm had stabilised. Recombinant proteins were eluted at 1 mL/min from the column into 5 mL fractions using the appropriate low salt buffer containing 300 mM imidazole. To cleave the N-terminal 6×His-SUMO tag, those fractions containing the protein of interest were pooled and dialysed overnight at 4°C into a cleavage buffer (50 mM HEPES-KOH (pH 7.4), 5 mM MgCl_2_, 2 mM BME, 20 mM imidazole, 100 mM KCl, 10% glycerol) in the presence of the SUMO protease, 6×His-tagged ULP1 (4 µg/mL per mg of recombinant protein). Cleaved proteins were loaded back onto the HisTrap^™^ HP Sepharose column under the same conditions as described previously to separate the 6×His-SUMO tag. The proteins were then loaded onto a HiLoad^®^ 16/600 Superdex^®^ 200 size-exclusion column (Cytiva) pre-equilibrated in 50 mM HEPES-KOH (pH 7.4), 5 mM MgCl_2_, 2 mM BME, 10% glycerol and 500 mM (DNAJB1) or 100 mM (HspA8 and Hsp110) KCl. Proteins were eluted from the column at 1.5 mL/min into 10 mL fractions. Fractions containing the protein of interest were pooled and concentrated using a 10K MWCO Pierce^™^ Protein Concentrator and stored at −80°C until use.

### 5.3 Preparation of α-synuclein seeds and mature fibrils

To form α-synuclein amyloid seeds, 100 μM monomeric α-synuclein was incubated in 50 mM phosphate buffer (PB; pH 6.3) in low protein binding tubes (Thermo Fisher Scientific) for 24 h at 45°C with constant stirring using a Spinbar® magnetic stirring flea (Sigma-Aldrich) on a Magnetic Stirrer & Hot Plate combination with ‘Thermostat’ temperature control (Industrial Equipment and Control, Thornbury, VIC, Australia). Samples were probe sonicated (Digital Sonifier 250, Branson Ultrasonics Corporation) at 30% power for 30 s (10 s on/30 s off) and were then incubated as before for a further 24 h. The sample was centrifuged (20,000 × *g*, 40 min, 4°C) to separate fibrils (which form a pellet) from soluble monomeric protein. A wash step was performed whereby the pellet was gently resuspended in 50 mM PB (pH 7.4) and centrifuged again (20,000 × *g,* 15 min, 4°C). Once the soluble fraction was removed and discarded, the pellet was resuspended in 50 mM PB (pH 7.4). The fibrils were sonicated a final time, as before, to fragment them into seeds. The final concentration of α-synuclein in these samples, reported as monomeric equivalent of α-synuclein, was determined via a BCA assay and aliquots were frozen in liquid nitrogen and stored at −80°C.

Seeds of α-synuclein (2.5 µM of monomeric equivalent) were elongated into mature fibrils by the addition of 100 µM of monomeric α-synuclein in 50 mM PB (pH 7.4) and incubation for 48 h at 45°C without shaking. Following the incubation period, samples were centrifuged (20,000 × *g,* 40 min, 4°C) to separate the mature fibrils from monomeric and smaller oligomeric forms of α-synuclein. A wash step was performed whereby the pellet was gently resuspended in 50 mM PB (pH 7.4) and centrifuged again (20,000 × *g,* 15 min, 4°C). Once the soluble fraction was removed and discarded, the pellet containing the mature fibrils was resuspended in 50 mM PB (pH 7.4) and stored at 4°C.

### 5.4 Assay to monitor the disaggregation of fibrillar forms of α-synuclein by molecular chaperones

To monitor the kinetics of the disaggregation of α-synuclein seeds, 2 µM α-synuclein seeds (monomeric equivalent) were incubated with 2 µM HspA8, 1 µM DNAJB1, and 0.2 µM Hsp110 in disaggregation buffer consisting of 50 mM HEPES-KOH, 50 mM KCl, 5 mM MgCl_2_, 2 mM DTT, 0.05% (v/v) Tween-20 (pH 7.5) supplemented with 20 µM Thioflavin-T (ThT). Samples were prepared in duplicate and incubated in a clear-bottom 384-well microplate (Greiner Bio-One, Kremsmünster, Austria) at 30°C in a FLUOstar Optima plate reader (BMG LABTECH, Melbourne, VIC, Australia) for 15 min for temperature equilibration of the ThT. Following this, pre-heated ATP (or Milli-Q® water as a control) was added to all samples (5 mM final ATP concentration) and the plate was shaken for 30 s. The subsequent disaggregation of α-synuclein fibrils was monitored by measuring the fluorescence of ThT using excitation/emission filters of 440/490 nm respectively, for a maximum of 16 h.

To investigate the effect of sHsps on the capacity of the Hsp70 chaperone system to disaggregate α-synuclein seeds, the seeds were first pre-incubated with sHsps (αB-crystallin or Hsp27) or ovalbumin (as a non-chaperone control protein) at molar ratios of 10:1, 2:1, 1:1, 1:2 or 1:10 (α-synuclein:sHsp/ovalbumin) for 30 min at room temperature to facilitate binding (Cox et al. 2018). Following this incubation, samples were diluted into disaggregation buffer containing 2 µM HspA8, 1 µM DNAJB1, 0.2 µM Hsp110 and 20 µM ThT in a clear-bottom 384-well black microplate (Greiner Bio-One) and incubated for 15 min prior to the addition of pre-heated 5 mM ATP. The ThT fluorescence of these samples was monitored as described above. For disaggregation experiments performed in the presence of monomeric α-synuclein, 10 µM monomeric α-synuclein was added to the disaggregation buffer along with the Hsp70 chaperone system.

For disaggregation experiments performed at physiological concentrations of α-synuclein and molecular chaperones, the following concentrations were used: 50 µM for monomeric α-synuclein (Iwai et al. 1995, Wilhelm et al. 2014); 20 µM for the sHsps (Mymrikov et al. 2020); 14 µM for HspA8 (Moran Luengo et al. 2018) with DNAJB1 and Hsp110 concentrations of 7 µM and 1.4 µM respectively to maintain the 1:0.5:0.1 molar ratio of HspA8:DNAJB1:Hsp110 used in previous experiments.

### 5.5 Assay to monitor the elongation of α-synuclein seeds in the presence of molecular chaperones

Samples of α-synuclein seeds (2 µM of monomeric equivalent) were incubated with up to 20 µM monomeric α-synuclein, 2 µM sHsp (or superoxide dismutase 1 (SOD1) as a non-chaperone control protein), 2 µM HspA8, 1 µM DNAJB1, and 0.2 µM Hsp110. As described previously, samples were prepared in duplicate in disaggregation buffer supplemented with 20 µM ThT and ThT fluorescence measured for up to 20 h at 30°C using a FLUOstar Optima plate reader (BMG LABTECH).

### 5.6 Native-PAGE and immunoblotting

To monitor the disaggregation of α-synuclein seeds by the Hsp70 chaperone system via Native-PAGE, 8 µM α-synuclein seeds (monomeric equivalent) were incubated with 8 µM HspA8, 4 µM DNAJB1, and 0.4 µM Hsp110 and 5 mM ATP in disaggregation buffer in low protein binding tubes (Thermo Fisher Scientific) and incubated at 30°C for a maximum of 4 h. Aliquots (50 µL) were taken from this mixture with 50 mM EDTA added (to a final volume of 55 µL) to quench the reaction for 20 min at room temperature before being placed on ice. Samples containing α-synuclein seeds only, monomeric α-synuclein only or α-synuclein seeds with chaperones in the absence of ATP were included as controls. Samples were subsequently diluted into Native-PAGE loading buffer (final concentrations: 0.81 M Tris-HCl (pH 6.8), 16% (v/v) glycerol, 0.01% bromophenol blue) before being loaded into the wells of a hand-cast 6% (w/v) polyacrylamide gel (4.7 mL Milli-Q® water, 3.7 mL Tris (1 M, pH 8.8), 1.5 mL 40% (w/v) bis-acrylamide (37.5:1), 25 µL 10% (w/v) ammonium persulfate, 25 µL tetramethylethylenediamine). Samples were subject to electrophoresis using a Mini-Protean® Tetra Cell system (Bio-Rad, Hercules, CA, USA) for 2.5 h at 150 V in ice-cold Native-PAGE running buffer (0.25 M Tris, 1.92 M glycine, pH 8.8).

Following Native-PAGE, samples were transferred to a PVDF membrane at 100 V in ice-cold transfer buffer (25 mM Tris base, 192 mM glycine, 20% (v/v) methanol, pH 8.3). The membrane was blocked overnight at 4°C in Tris-buffered saline (TBS; 50 mM Tris base, 150 mM NaCl, pH 7.6) containing 5% (w/v) skim milk powder. Membranes were then probed with an anti-α-synuclein antibody (mouse monoclonal, Abcam, Cambridge, UK, catalogue number ab1903) diluted 1:2000 in TBS supplemented with 0.05% (v/v) Tween-20 (TBS-T) containing 5% (w/v) skim milk powder. Primary antibody incubation was performed for 2 h at room temperature with gentle rocking. The membrane was washed four times (each for 10 min) in TBS-T before being incubated with the secondary antibody (anti-mouse IgG HRP-conjugated antibody; rabbit polyclonal, Sigma-Aldrich; #A9044) diluted 1:5000 in TBS-T containing 5% (w/v) skim milk powder for 1 h with gentle rocking at room temperature. Finally, the membrane was washed as before in TBS-T before visualisation of proteins of interest with SuperSignal® West Dura Extended Duration Chemiluminescent Substrate (Thermo Fisher Scientific) using a ChemiDoc™ Imaging System (Bio-Rad).

### 5.7 ThT fluorescence data analysis

For each sample, background ThT fluorescence of buffer and chaperones was subtracted. At each time point, the fluorescence intensity of samples was calculated relative to the fluorescence intensity of samples containing α-synuclein fibrils or seeds alone. Hence, the normalised change in ThT fluorescence reflects the ThT fluorescence intensity at any given timepoint as a percentage of the fibril/seed-only control. Data from the technical duplicates were averaged and, where relevant, each sample was fit to a one-phase exponential decay model in R (R Core Team 2022). For each sample, the rate and endpoint of the fitted model were extracted to quantify the rate of disaggregation and disaggregation yield, respectively.

At the end of each experiment involving addition of monomeric α-synuclein, the ability of each chaperone mixture to inhibit α-synuclein aggregation was calculated using the following formula: Protection (%) = 100 × (ΔF_N_ – ΔF_C_)/ΔF_N_ where ΔF_C_ and ΔF_N_ represent the normalised overall change in ThT fluorescence in the presence and absence of chaperones, respectively.

Where appropriate, to test for synergism or antagonism, the fractional effect of each chaperone class (sHsp or Hsp70 system) in reducing α-synuclein seed aggregation was calculated using:

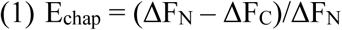

where ΔF_C_ and ΔF_N_ represent the normalised overall change in ThT fluorescence in the presence and absence of chaperones, respectively. Then, the expected additive effect (E_sHsp + Hsp70_) of combining these two chaperone classes was calculated in accordance with the Bliss independence model (Bliss 1939) using:

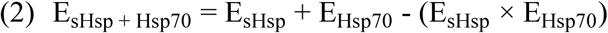

where E_sHsp_ and E_Hsp70_ refer to the fractional effect of either sHsp or the Hsp70 chaperone system and E_sHsp + Hsp70_ refers to the expected additive effect of the combination. If the measured fractional effect was greater than the E_sHsp + Hsp70_, the combination of chaperones was considered synergistic and if the measured fractional effect was less than E_sHsp + Hsp70_ the combination of chaperones was considered antagonistic.

### 5.8 TIRF Microscopy

Samples were imaged using a custom-built TIRF microscopy setup as previously detailed (Marzano et al. 2022). To prepare coverslips for imaging, 24 mm by 24 mm glass coverslips were cleaned by alternate cycles of water bath sonication (Digital benchtop ultrasonic cleaner 250HD, Soniclean, Dudley Park, SA, Australia) for 20 min in 100% ethanol and KOH (5 M), repeated three times. A final sonication in Milli-Q® water for 15 min was performed.

Coverslips were dried with compressed nitrogen gas and incubated with poly-L-lysine solution (0.01% (w/v)) for 30 min to immobilise α-synuclein fibrils to the surface. Following this incubation, coverslips were rinsed with Milli-Q® water, dried using compressed nitrogen gas and stored in a custom humidity chamber before use. Coverslips were coupled to the objective lens by immersion oil before loading of the sample.

All α-synuclein fibrils and seeds were diluted approximately 1:500 in 50 mM PB (pH 7.4) containing 6 mM 6-hydroxy-2,5,7,8-tetramethyl chroman-2-carboxylic acid (TROLOX) and an α-cyanostilbene derivative (ASCP) dye (5 μM) (Marzano et al. 2020). Samples (50 μL) were deposited directly onto the surface of the coverslip and illuminated using a circularly polarised laser (Sapphire 488-150 CW, Coherent, Saxonburg, PA, USA) at 200 mW cm^-2^ at 488 nm. The camera (Andor iXon Life 897 EMCCD, Oxford Instruments, Abingdon, UK) was operated at −70°C in frame transfer mode at 20 Hz with an electron multiplication gain of 700 and pixel distance of 160 nm (in sample space). Imaging of the ASCP fluorescence was performed at 500 ms per image. Samples were at room temperature during imaging (approx. 20°C).

All TIRF images were initially corrected for electronic offset and uneven excitation beam distribution across the field of view. To determine the length of α-synuclein seed and fibril contours, corrected images were analysed using the “Ridge Detection” plugin (Steger 1998) in Fiji ImageJ (Schindelin et al. 2012), providing the length of seed or fibril contours (in pixels). These lengths were converted to nm and deconvolved using the following formulae:

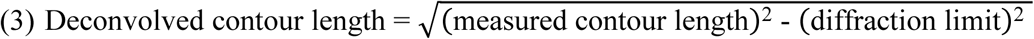

Where the diffraction limit is calculated as

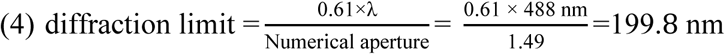

### 5.9 Statistics

All statistical analyses were performed in R (R Core Team 2022), using ggbreak (Xu et al. 2021) (version 0.1.2), outliers (Komsta 2022) (version 0.15), emmeans (Lenth 2023) (version 1.8.9) and car (Fox and Weisberg 2019) (version 3.1-1) packages. Data were assessed for normality via the Shapiro-Wilk test and homoscedasticity via Cochran’s C test. Where necessary, the overall change in ThT fluorescence (endpoint) data were log_10_ or square-root transformed to improve normality and homoscedasticity. A one-way analysis of variance (ANOVA) with Tukey’s honestly significant difference (HSD) post-hoc test was performed to determine statistically significant differences between groups in overall change in ThT fluorescence, rate of disaggregation or overall amount of disaggregation. A P-value of less than 0.05 was considered statistically significant.

## 6 AUTHOR CONTRIBUTIONS

Nicola Auld: Conceptualization; writing – original draft; writing – review and editing; visualization; formal analysis; data curation; investigation. Shannon McMahon: writing – review and editing; resources. Nicholas Marzano: Writing – review and editing; supervision. Antoine van Oijen: Conceptualization; Writing – review and editing; supervision; funding acquisition. Heath Ecroyd: Conceptualization; writing – review and editing; project administration; supervision; funding acquisition.

## 7 ACKNOWLEDGEMENTS

We thank the University of Wollongong staff that provided technical and administrative support for this work. This work was funded by the Australian Research Council (DP220103466). AMvO also acknowledges funding from the National Health and Medical Research Council (APP1197069).

