## Supplementary for "Competing chaperone pathways in α-synuclein disaggregation and aggregation dynamics"

### SUPPLEMENTARY MATERIAL

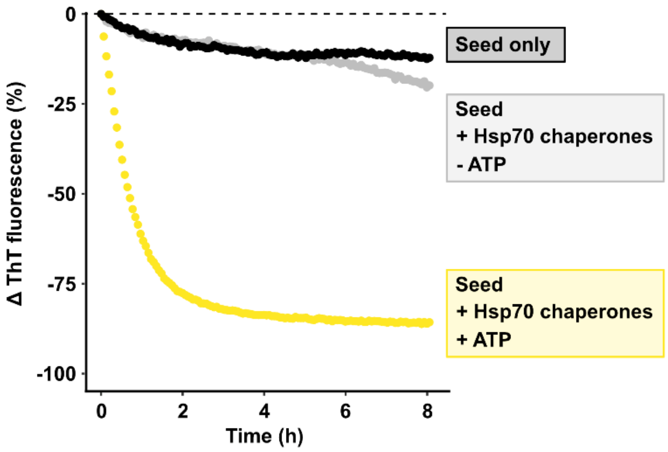


**Supplementary Figure 1. Hsp70-mediated disaggregation is ATP dependent.** α-synuclein seeds (2 µM) were incubated with Hsp70 chaperones (2 µM HspA8, 1 µM DNAJB1 and 0.2 µM Hsp110) in the presence or absence of ATP at 30ºC for up to 8 h. Kinetic traces of α-synuclein seed and fibril disaggregation as monitored by the change in ThT fluorescence over time. Data are representative of three independent experiments.

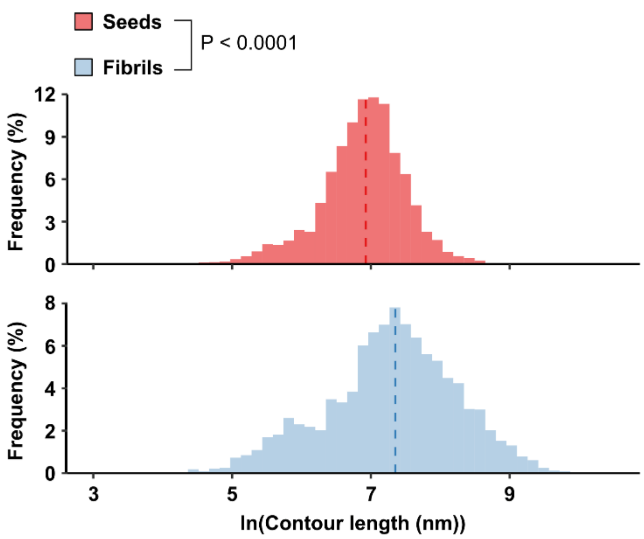


**Supplementary Figure 2. Sonication of mature α-synuclein fibrils produces shorter amyloid fragments (seeds).** Lengths of mature fibrils and α-synuclein seeds were calculated from TIRF microscopy images using the Ridge Detection plugin (Steger 1998) in ImageJ Fiji. Data were analysed using a two-tailed Welch's t-test (n = 8609 identified contours for seeds and n = 3303 identified contours for fibrils taken from at least 30 images of each treatment from 2 independent experiments).
